# Structural basis of sRNA RsmZ regulation of *Pseudomonas aeruginosa* virulence

**DOI:** 10.1101/2022.11.16.516715

**Authors:** Xinyu Jia, Zhiling Pan, Yang Yuan, Bingnan Luo, Yongbo Luo, Sunandan Mukherjee, Guowen Jia, Liu Liu, Xiaobin Ling, Xiting Yang, Zhichao Miao, Xiawei Wei, Janusz M. Bujnicki, Kelei Zhao, Zhaoming Su

**Author notes:** Correspondence should be addressed to Z.S. and K.Z. These authors contributed equally (Xinyu Jia, Zhiling Pan, Yang Yuan).

## Abstract

*Pseudomonas aeruginosa* is an opportunistic human pathogen that poses threats to hospitalized immunocompromised patients ^1^. A non-coding small RNA (sRNA) from the repressor of secondary metabolites (Rsm) system, RsmZ, sequesters the global repressor protein RsmA to regulate downstream gene expressions that reprogram virulence repertoires associated with acute and chronic *P. aeruginosa* infections ^2,3^. Molecular insights into the full-length RsmZ architecture remain elusive, leading to the lack of understanding of RsmZ binding to RsmA and subsequent modulations of gene expressions. Here we use cryo-electron microscopy (cryo-EM) to resolve structures of the full-length RsmZ in complexes with RsmA, in which five stem-loops (SLs) and one single-stranded junction carrying the GGA binding sites in RsmZ form three pairs of clamps, each binding to a RsmA homodimer. Disruptions of the base-pairings in all stems of RsmZ significantly reduced the binding affinity to RsmA by 17-fold, which resulted in enhanced RsmA downregulation of gene expressions and phenotypes associated to both acute and chronic virulence of *P. aeruginosa*. Double mutations that rescued these stems of RsmZ restored the binding to RsmA by more than 5-fold, and recovered the corresponding phenotypes. Our results reveal the molecular mechanism of RsmZ regulation of *P. aeruginosa* virulence and suggest RsmZ as a potential target for the development of new antimicrobial agents.

## Introduction

*Pseudomonas aeruginosa* is a ubiquitous Gram-negative opportunistic bacterium that notoriously causes infections with high mortality rate in hospitalized patients, especially those with compromised immune systems ^4^. *P. aeruginosa* can cause acute infections that are typically associated with the cytotoxins secreted by the type III secretion system (T3SS) ^5^, as well as chronic persistence relied on T6SS and biofilm formation, including those in cystic fibrosis patients ^6^. An intercellular communication network based on cell density, quorum sensing (QS), regulates numerous gene expressions including those related to both acute and chronic virulence of *P. aeruginosa* ^7^. The global regulatory protein of the repressor of secondary metabolites (Rsm) system in bacteria, RsmA, is found to predominantly regulate these virulence entities in *P. aeruginosa* by modulating related gene expressions at transcription and posttranscription levels ^8-13^.

A non-coding small RNA (sRNA) RsmZ, whose expression is regulated by the GacS/GacA two-component system ^14^, can sequester RsmA and antagonize its repressive function, thereby promoting downstream gene expressions associated with *P. aeruginosa* virulence ^2,3^. Previous structural and functional studies in *Pseudomonas* genus have revealed how homodimers of RsmA and other Rsm family member proteins recognized the GGA binding sites on two separate stems that formed a clamp shape ^2,15-18^. In particular, two conformations in solution of the truncated RsmZ with the first four stems in complexes with three RsmE homodimers in *P. fluorescens* were derived using a combination of nuclear magnetic resonance (NMR) and electron paramagnetic resonance (EPR) spectroscopy ^15^. Although that work revealed a cooperative mechanism of RsmE assembly on RsmZ, the architecture of full-length RsmZ and molecular mechanism of RsmZ sequestration of RsmA in the pathogenic *P. aeruginosa* remains elusive, which continues to limit our understanding of RsmZ regulation of *P. aeruginosa* virulence.

Single particle cryo-electron microscopy (cryo-EM) can resolve structures in different conformations of biological macromolecules at near-atomic resolution ^19,20^. Here we use cryo-EM to determine structures of the full-length RsmZ in complexes with RsmA homodimers in *P. aeruginosa*. Five stem-loops (SL1-SL5) and the single-stranded junction between SL2 and SL3 (J2/3) carrying the GGA binding sites form three pairs of clamps, each binding to a RsmA homodimer. The terminator SL (SL_ter_) coaxially stacks on SL5 and points away from the rest of the complex. Another conformation with one missing RsmA homodimer between SL2 and SL3 reveals the dynamics of SL_ter_ upon RsmA binding. Previous study indicated that the base-pairing stem affects RsmA binding ^16^. Mutations that disrupt all five stems in RsmZ lead to 17-fold decreased binding affinity to RsmA and downregulation of gene expressions associated to QS, T6SS and biofilm, leading to reduced phenotypes of proteolytic activity, pyocyanin generation and biofilm formation in *P. aeruginosa*. Double mutations that rescue these stems of RsmZ recover the binding to RsmA, the downstream gene expression levels and the corresponding phenotypes. These results elucidate the structural basis of full-length RsmZ binding to RsmA, and provide molecular insights into RsmZ regulation of *P. aeruginosa* virulence, suggesting RsmZ as a potential RNA target for novel antimicrobial development.

## Results

### Cryo-EM structures of RsmZ-A complexes

Incubation of separately purified full-length RsmZ from *in vitro* transcription and RsmA from recombinant expression in *Escherichia coli* resulted in ribonucleoprotein (RNP) complex formation when the ratio of protein to RNA reached 4 to 1, as observed in electrophoretic mobility shift assay (EMSA) (Extended Data Fig.1a). Additions of different detergents, including lauryl maltose neopentyl glycol (LMNG), octyl-beta-glucoside (β-OG) or glyco-diosgenin (GDN) facilitated more efficient complex formations in the presence of less protein, indicating that these detergents may enable faster formations of more stable RNP complexes (Extended Data Fig.1b-d).

Cryo-EM single particle analysis of the RsmZ-A complex with GDN yielded a three-dimensional (3D) reconstruction of the full-length RsmZ in complex with three RsmA homodimers (RsmZ-A_3_) at 3.80 Å resolution (Fig.1a, Extended Data Fig.2). RsmZ consists of six consecutive SLs, namely SL1-SL5 and a SL_ter_ toward the 3’ end stacking on SL5 and points away from the rest of the complex (Fig.1b-c), which is distinct from the previously predicted secondary structure ^2^. Six GGA binding sites in the loop regions of SL1-SL5 and a single-stranded junction J2/3 are grouped into three pairs, SL1 and SL5, SL2 and SL3, J2/3 and SL4, with each resembling a clamp shape that binds a RsmA homodimer via interactions between the GGA motifs of RsmZ and residues 36-44 of RsmA (Fig.1c, Extended Data Fig.3a-f), similar to previous truncated structures of homologous RsmZ-E in *P. fluorescens* and RsmZ-N in *P. aeruginosa* (Extended Data Fig.3g-i) ^15,18^. Previous study has derived structures of truncated RsmZ with SL1-SL4 (nt 1-72) binding to RsmE in *P. fluorescens* using NMR and EPR ^15^. Sequence alignment and covariance analysis among *Pseudomonas* RsmZs revealed that SL1-SL4 and SL_ter_ are structurally conserved between *P. aeruginosa* and *P. fluorescens* (Extended Data Fig.4). However, comparisons of the full-length and truncated RsmZs reveal substantially different architectures, with only one of the RsmA binding clamps, consisting of SL2 and SL3 in both RsmZs, interacting with RsmA homodimer in a similar way (Extended Data Fig.5).

**Fig.1.**
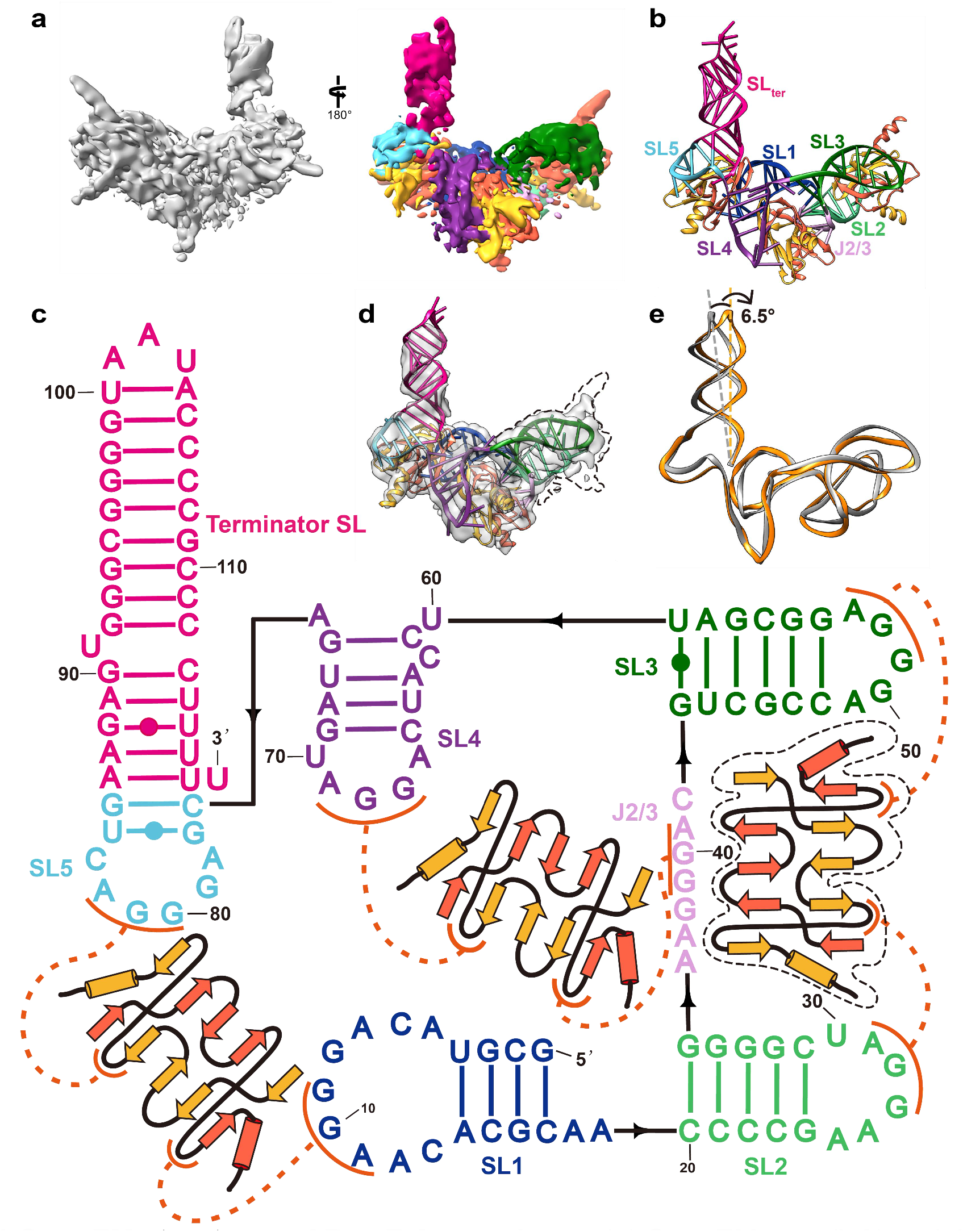
Cryo-EM structures of RsmZ-A complexes. (**a**) Cryo-EM map (left) and the segmented map (right) of RsmZ in complex with three RsmA homodimers coloured according to the secondary structure. (**b**) Coloured cryo-EM model of RsmZ in complex with three RsmA homodimers. (**c**) Secondary structure of the RsmZ-A cryo-EM model. (**d**) Coloured cryo-EM map and model of RsmZ in complex with two RsmA homodimers. (**e**) Overlap of the cryo-EM structures of RsmZ in complexes with two (orange) and three RsmA homodimers (gray).

Another state of the RsmZ-A complex with one RsmA homodimer missing between SL2 and SL3 was isolated from 3D classification (Fig.1d). The superposition of both RNP structures reveals a ∼6.5º rotation of SL_ter_ (Fig.1e), indicating the dynamics of SL_ter_ during RsmA binding, which is also supported by the local resolution map (Extended Data Fig.2b, c).

### Binding of RsmZ to RsmA homodimers

Previous studies observed sequential and cooperative bindings of RsmE to RsmZ in *P. fluorescens*, in which binding affinity was affected by the sequence of the loop and closing base pairs of the stem ^15,16^. In contrast, we observed simultaneous binding of all three RsmA homodimers to RsmZ in EMSA (Extended Data Fig.1). To evaluate the critical roles that RsmZ structure plays in RsmA binding in *P. aeruginosa*, we first determined the dissociation constant (K_D_) of RsmZ binding to RsmA as 194±15 nM by surface plasma resonance (SPR) (Fig.2a). Mutations that disrupted all five stems (RsmZ_mut-all_) led to the K_D_ of 3350±560 nM, decreased by 17-fold (Fig.2b), whereas double mutations that recovered all stems (RsmZ_recov-all_) bound to RsmA with the restored K_D_ of 634±111 nM (Fig.2c). Individual K_D_ for each binding site to one RsmA homodimer upon mutations of the rest of the GGA sequences was determined as 385±38 nM for SL1 and SL5, 319±27 nM for SL2 and SL3, and 789±20 nM for J2/3 and SL4, respectively (Fig.2a, Extended Data Fig.6a-c). The overall comparable K_D_ of each binding site indicates a simultaneous binding mechanism of RsmA to RsmZ in *P. aeruginosa*.

**Fig.2.**
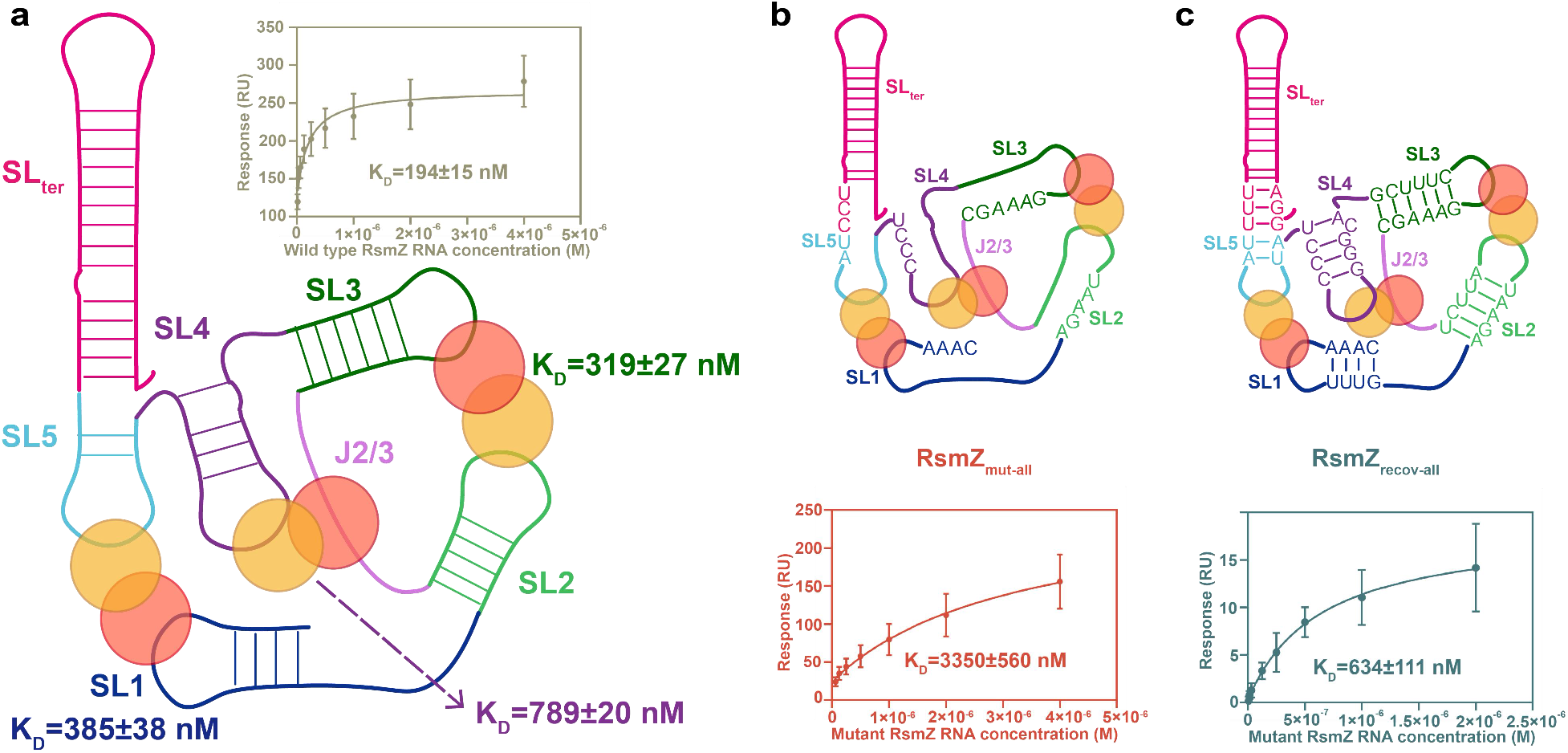
Binding affinity of RsmA to WT and mutant RsmZ, and individual binding sites of RsmZ. (**a**) Binding affinity of RsmA to WT RsmZ and individual binding site. (**b**) Binding affinity of RsmA to RsmZ_mut-all_. (**c**) Binding affinity of RsmA to RsmZ_recov-all_. Sites of mutations introduced to WT RsmZ are shown.

### RsmZ regulation of *P. aeruginosa* virulence

Non-coding sRNAs RsmY and RsmZ bind to RsmA to regulate downstream gene expression related to *P. aeruginosa* virulence ^21^. Compared to wild-type (WT) *P. aeruginosa* reference isolate PAO1, RsmY and RsmZ double-knockout mutant (ΔRsmYZ) showed decreased expression levels of *pslA* that controls biofilm formation, *lasB, rhlA* and *phzS* that characterize QS activity, and *hcp1* associated to H1-T6SS, leading to reduced biofilm formation and production of QS-controlled extracellular proteases and pyocyanin (Fig.3). Expression of WT full-length RsmZ in PAO1_ΔRsmYZ_restored the gene expressions and the corresponding phenotypes back to the levels comparable to those of WT PAO1, whereas expression of RsmZ_mut-all_with disrupted base-pairings in SL1-SL5 of RsmZ in PAO1_ΔRsmYZ_did not significantly change gene expressions and phenotypes of PAO1_ΔRsmYZ_, due to the diminished binding of RsmZ_mut-all_to RsmA. Expression of RsmZ_recov-all_with double mutations that recovered base-pairings of SL1-SL5 of RsmZ in PAO1_ΔRsmYZ_recovered the gene expressions and phenotypes comparable to those of WT PAO1, consistent with the restored binding affinity of RsmZ_recov-all_to RsmA (Fig.3). This suggests a RsmZ regulation mechanism through RsmA binding, and demonstrates the critical role that RsmZ structure plays in regulation of *P. aeruginosa* virulence.

**Fig.3.**
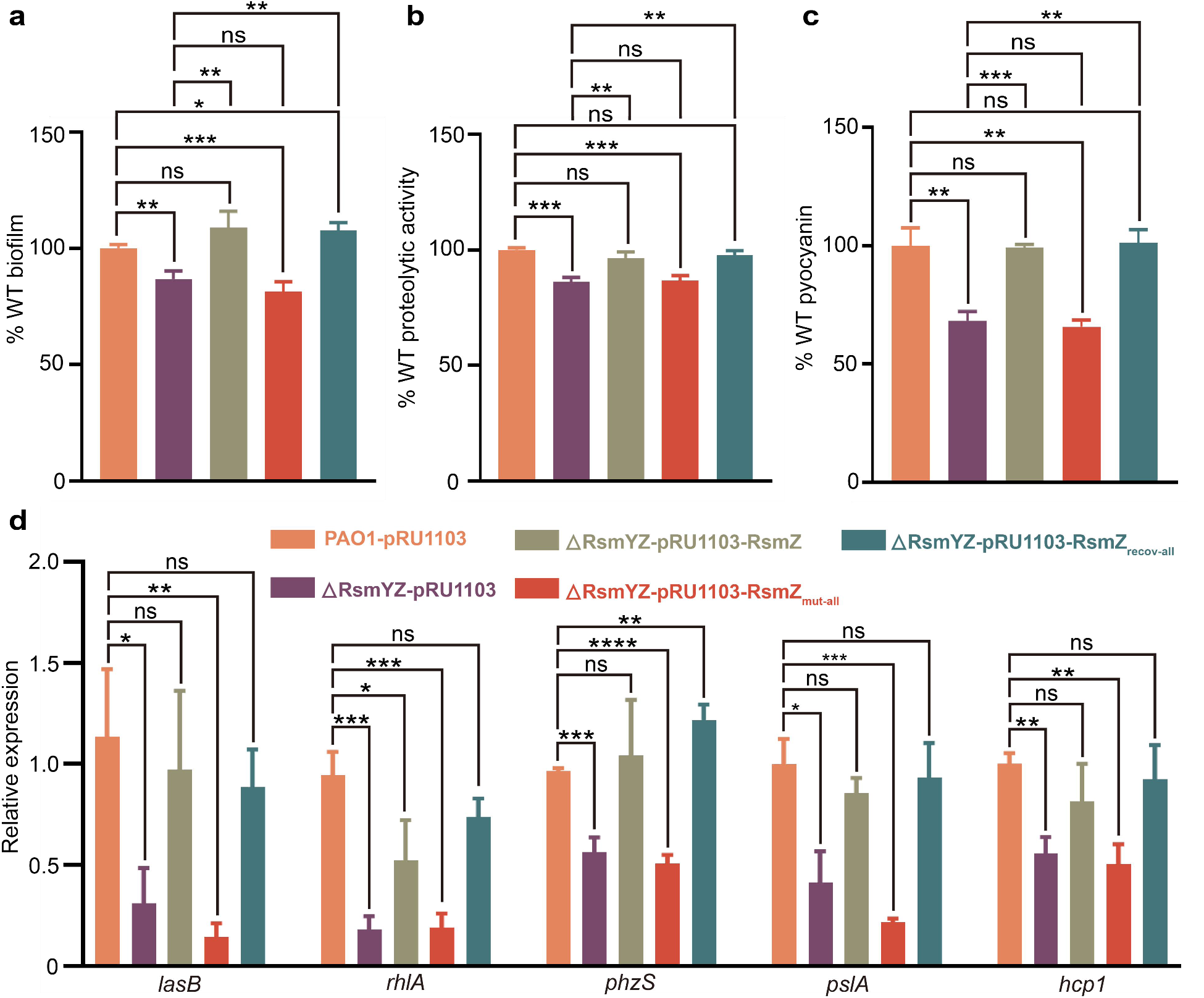
Regulation of WT and mutant RsmZ to gene expressions and phenotypes associated to *P*.*aeruginosa* virulence. (**a**) Regulation of RsmZ on *P*.*aeruginosa* biofilm formation. Biofilm was measured by crystal violet staining and values were normalized to wild-type WT PAO1. (**b**) Regulation of RsmZ on *P*.*aeruginosa* proteolysis. Diameter values of proteolytic rings in the M9-skim milk plate were normalized to WT PAO1. (**c**) Regulation of RsmZ on *P*.*aeruginosa* pyocyanin production. Pyocyanin was detected after culture in LB medium and normalized to WT PAO1. (**d**) Elastase (*lasB*), rhamnolipid production (*rhlA*), pyocyanin (*phzS*), biofilm (*pslA*) and H1-T6SS (*hcp1*) gene expression were determined by qRT-PCR. Data shown are means ± SD of three independent replicates. One-way ANOVA. ns, *P*>0.05. *, *P*<0.05. **, *P*<0.01. ***, *P*<0.001. ****, *P*<0.0001. WT PAO1 with empty pRU1103 was in orange. RsmYZ knock out PAO1 with empty pRU1103 was in purple. RsmYZ knock out PAO1 with different RsmZ constructs were colored the same as the binding curves in Figure2.

## Discussion

Bacterial sRNAs participate in various important biological processes such as transcription, translation, and catalysis ^22^. RsmZ in *P. aeruginosa* has been found to regulate gene expressions by sequestration of the RsmA protein, a homolog of carbon storage regulator A (CsrA) that globally regulates gene expression levels related to metabolism, QS, and virulence ^21^. Although NMR combined with EPR provided some structural insights of truncated RsmZ binding to RsmE in *P. fluorescens* ^15^, the full-length RsmZ-A complex structure in *P. aeruginosa* remains unknown, leading to the paucity in understanding of the essential role that RsmZ structure plays in RsmA sequestration and gene expression regulations.

Recent advances in single particle cryo-EM allowed structure determinations of RNAs and RNPs ^19^. We resolved cryo-EM structures of the full-length RsmZ in complex with two or three RsmA homodimers, and revealed a distinct architecture of RsmZ compared to either conformation of the truncated RsmZ in *P. fluorescens* (Extended Data Fig.5). Binding experiments suggested simultaneous binding of RsmA to RsmZ, whereas disruptions of stems enclosing each binding site significantly decreased binding affinity to RsmA. Subsequent functional experiments demonstrated that maintaining the intact RsmZ architecture is crucial for regulation of QS, T6SS and biofilm formation associated to *P. aeruginosa* virulence (Fig.4). The overall conserved sequences and structures of *Pseudomonas* RsmZ suggests a universal RsmA sequestration mechanism regulated by the RsmZ structure. Together, our results provide the structural basis of RsmA sequestration by RsmZ, and establish the critical role of sRNA architecture in regulations of *P. aeruginosa* virulence, which may facilitate novel RNA-targeted antimicrobial development.

**Fig.4.**
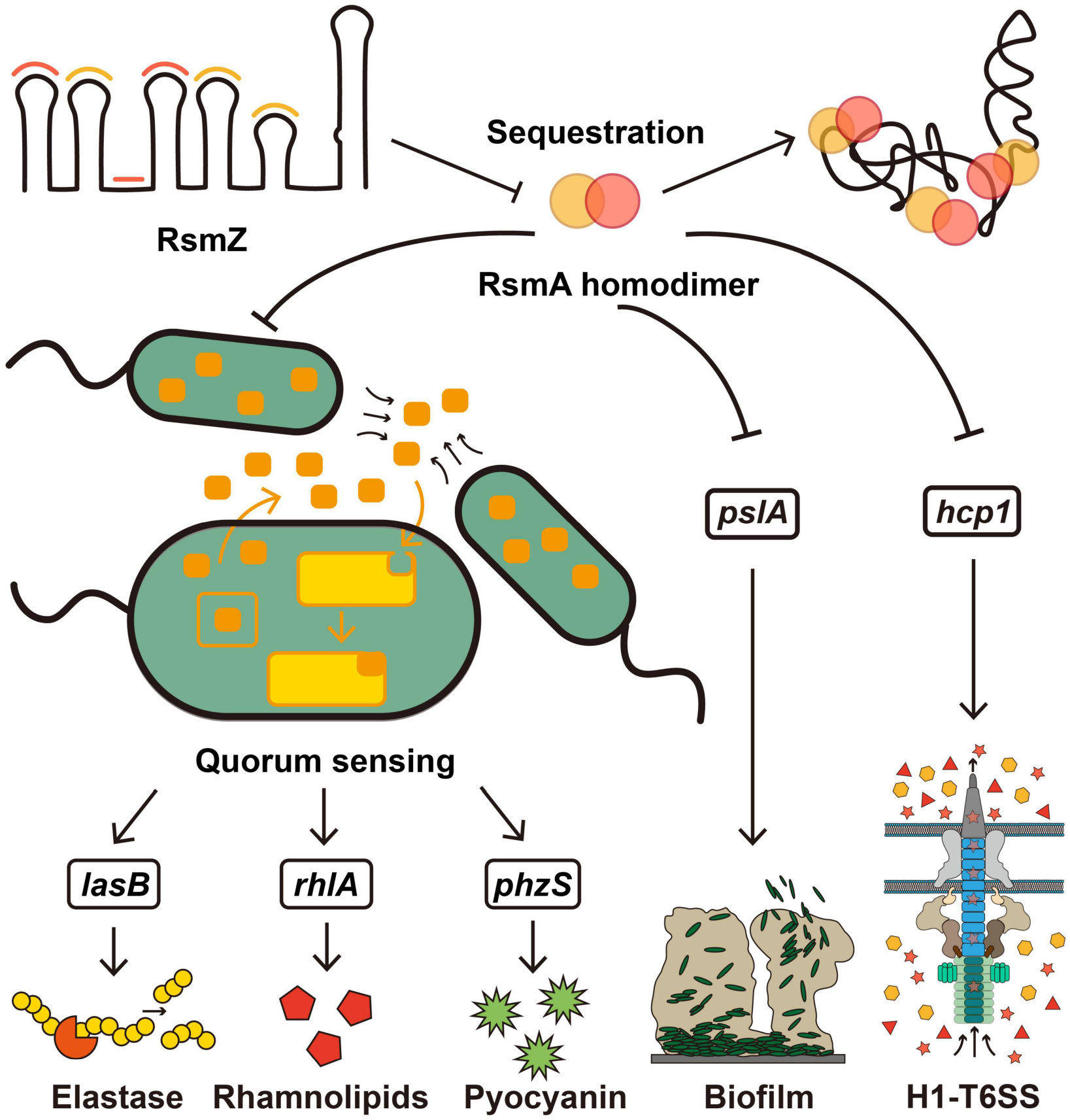
Schematic representation of RsmZ regulation of *P*.*aeruginosa* virulence.

## Materials and Methods

### RsmA protein expression and purification

The pET28a expression vector encoding His-tagged RsmA was transformed into *E. coli* BL-21 (DE3). The transformed cells were grown in LB medium at 37°C to OD600 = 0.6-0.8 before induction with 0.5 mM IPTG and then cultured at 18°C for 16 h. Cells were pelleted and lysed in 20 mM Tris-HCl, pH=7.5, and 150 mM NaCl (Buffer A) via sonication. The lysate was centrifuged, and the pellet was washed with lysis buffer before it was solubilized in 20 mM Tris-HCl, pH=7.5, 150 mM NaCl (Buffer B). Clarified lysate was filtered and was initially purified on His-select resin equilibrated in the lysis buffer and washed with lysis buffer containing 20 mM imidazole, and eluted with lysis buffer containing 500 mM imidazole. Protein samples were concentrated and further purified by size exclusion chromatography on a Superdex 75 column (GE Healthcare Life Sciences) pre-equilibrated with 20 mM Tris-HCl, pH=7.5, 150 mM sodium chloride, and 2 mM TCEP.

### RNA preparations

RNA was prepared as previously described ^15^. Briefly, the DNA template was amplified from the pUC19-T7-template. The DNA templates of all RNAs with 3’-OH were amplified by PCR with reverse primers containing two 2’-O-Methyl modified bases at the 5’ end to yield homogeneous 3’-OH on the RNAs’ 3’ ends ^23^. The RNA was prepared by *in vitro* transcription in a reaction containing 0.2 μM DNA template, 40 mM Tris-HCl, pH=7.9, 20 mM MgCl_2_, 2 mM spermidine, 0.01% TritonX-100, 10 mM DTT, 4 mM NTPs, and 0.8 μg/μL T7 RNA polymerase. The transcription reaction was incubated at 37°C for 2-3 h. The RNA was then centrifuged at 13,000 rpm for 5 min and the supernatant was mixed with loading buffer containing 6M urea, 1×Tris/Borate/EDTA buffer, 0.1% xylene cyanol, and 0.1% bromophenol blue and loaded on an 8% 29:1 acrylamide:bis, 8 M urea polyacrylamide gel. The gel electrophoresis was performed at 300V for 2-3 h, then visualized briefly with a 254-nm UV lamp, held far from the gel to minimize RNA damage ^24^. RNA was eluted from the gel overnight in 30 mM NaOAc pH=5.2, 0.1 mM EDTA at 4°C, then concentrated with Amicon® Ultra-15 10K Centrifugal Filter Devices (Millipore) to a total volume of 500 μL and precipitated with isopropanol.

### RNA biotinylation

Freshly prepared 0.5 M NaIO_4_(10 µL) and 0.5 M NaOAc pH=4.7 (10 µL) were added to RNA (up to 1 nmol) to a final volume of 50 µL and the solution incubated at room temperature for 1.5 h. The excess of NaIO_4_was reduced with 0.25 M KCl (final concentration) for 10 min on the ice. The RNA was precipitated with isopropanol and diluted with 10 µL water. Then 5 µL 0.5 M NaOAc, pH=4.7, and 20 mM biotin-hydrazide (TargetMol) in DMSO were added. Coupling was carried out at room temperature overnight. The mixture was extracted with water saturated phenol. The RNA was precipitated with isopropanol.

### Surface plasmon resonance (SPR)

The experiments were conducted via Biacore X-100 and streptavidin (SA) sensor chips (BIAcore International SA, Neuchâtel, Switzerland). Biotinylated RNAs were refolded by the method mentioned above. SA sensor chips were activated by 50 mM NaOH and 1 M NaCl for 1 min and 3 times, and 2 µg/mL refolded biotinylated RNAs were immobilized on the sensor surface by flowing into the chamber channel. The K_D_of RsmZ binding to RsmA was obtained from serial 2× dilution of RsmA starting from 4000 nM. The sensor was regenerated in 0.5% SDS after each measurement. The K_D_values with standard deviations were calculated via Biacore X100 Evaluation Software Version 2.0.2 from two parallel experiments.

### Construction of vectors for WT RsmZ and RsmZ mutants RNA preparation and expression in PAO1

All vectors for WT RsmZ and RsmZ mutants RNA preparation were generated from plasmid pUC19 carrying WT RsmZ DNA template sequences. For K_D_estimation of one RsmA homodimer binding, the other GGA binding sites were converted into CCT. To evaluate the effects of RsmZ stems on RsmA binding, plasmids with RsmZ mutations that disrupt all stems were generated (Fig.2). All pUC19-T7-template vectors were obtained by QuickChange method. The RsmZ PAO1 expression plasmid was constructed by adding PC promoter upstream of the DNA template sequences through PCR and then assembling into the HindIII and SalI restriction sites of pRU1103. The RsmZ stems mutants PAO1 expression plasmids were assembled into the HindIII and KpnI restriction sites of pRU1103.

### Electrophoresis mobility shift assay (EMSA)

All RNA and protein used for EMSA were estimated by the A280/A260 method of the NanoDrop spectrophotometer (Thermo Scientific). All reactions were carried out in binding buffer (10 mM Tris-HCl pH=7.4, 100 mM KCl, 2 mM MgCl_2_). Refolded RsmZ was prepared by heating to 95°C for 3 min in 10 mM Tris-HCl, pH=7.4 and 100 mM KCl and then adding MgCl_2_to a final concentration of 2 mM on ice. Reactions were mixed to a final concentration of 0.5 µM refolded RsmZ and 0-2 µM RsmA, and incubated at 37°C for 30 min. The reactions were mixed with loading buffer containing 10 mM Tris-HCl pH=7.4, 100 mM KCl, 10 mM MgCl_2_0.1% xylene cyanol, 0.1% bromophenol blue and then loaded onto a native 6% acrylamide gel and electrophoresed at 4°C for 40 min at 110V. Gels were stained by Sybr Gold (Invitrogen) and imaged by ChemDoc XRS+ (BioRad).

### Cryo-EM sample preparation

A total of 3 μL of mixture treated as described in EMSA was applied onto glow-discharged (45 s) 200-mesh R2/1 Quantifoil Au grids. The grids were blotted for 2.5 s in 100% humidity with no blotting offset and rapidly frozen in liquid ethane using a Vitrobot Mark IV (Thermo Fisher).

### Cryo-EM single particle data acquisition and data processing

The abovementioned frozen grids were loaded in Titan Krios (Thermo Fisher) operated at 300 kV, condenser lens aperture 50 μm, spot size 5, parallel beam with illumine area of 1.03 μm in diameter. Microscope magnification was at 165,000× (corresponding to a calibrated sampling of 0.85 Å per physical pixel). Movie stacks were collected automatically using EPU software on a K2 Bioquantum direct electron device equipped with an energy filter operated at 20 eV (Gatan), operating in counting mode at a recording rate of 5 raw frames per second and a total exposure time of 6 seconds, yielding 30 frames per stack, and a total dose of 59.7 e^-^/Å^2^. Two datasets were collected and a total of 17,800 movie stacks were collected with defocus values ranging from −1.2 to −1.7 μm. These movie stacks were motion corrected using Motioncor2 ^25^. After CTF correction by CTFFIND4 ^26^, then subjected to EMAN2.2 for neural network particle picking ^27^. A total of 8,008,685 particles were extracted in Relion3 ^28^. After two independent rounds of 2D classifications of two datasets, the best classes were selected by visual examination. Then, a total of 2,782,191 particles were subjected to cryoSPARC3.2.0 to build the initial model and then subjected to heterogeneous refinement to yield a major class including 817,964 particles. After the 3D auto-refine in Relion using 817,964 particles, the result was low pass to 20, 30, 40, 50 and 60 Å as references. 817,964 particles were subjected to heterogeneous refinement using the previously gained references and yielded RsmZ-A_2_and RsmZ-A_3_initial maps, which were erasered, masked and low-passed to 20, 40 and 60 Å. And then another round of 3D classification in Relion was processed using 817,964 particles and yielded two classes of particles with RsmZ-A_2_and RsmZ-A_3_features. After 2 rounds of 3D classification and 2D classification, two classes of particles were yielded. Finally, the class including 211,463 particles with RsmZ-A_2_features was subjected to cryoSPARC 3D Local refinement, CTF refinement, Bayesian polishing and post-processing to yield the final map for RsmZ-A_2_. Meanwhile, the class including 483,925 particles with RsmZ-A_3_features was subjected to cryoSPARC 3D Local refinement, CTF refinement, Bayesian polishing and post-processing to yield the final map for RsmZ-A_3_. The final 3D maps were displayed in UCSF Chimera ^29^.

### Cryo-EM model building and refinement

The initial RsmZ-A complex model was built with SimRNA ^30^, then manually adjusted and rebuilt with Coot as needed ^31^. The models were refined with Phenix.real_space_refine ^32^, yielding model–map correlation coefficient (CCmask) of 0.56 for RsmZ-RsmA_3_and 0.68 for RsmZ-RsmA_2_. The final model was validated by MolProbity ^33^. Secondary structure diagrams were manually prepared in Adobe Illustrator.

### Covariation analysis of RsmZ secondary structure and sequences alignment

We used the full length sequence of RsmZ and the secondary structure, derived from our cryo-EM structures, as the input of Infernal 1.1.4 software ^34^ to perform initial structure searching within the all non-redundant complete *Pseudomonas* genome from *Pseudomonas* genome database (accessed on 2022-06-16) ^35^. In detail, the sequence and secondary structure of RsmZ was represented in *stockholm* format, which was used to build covariance model by *cmbuild*. Then, *cmsearch* was used to fetch matched sequences from above-mentioned sequence database. Matched sequences were filtered so that the resulting sequences must contain >80% of the original structure length, and the redundant sequences were removed. A new *stockholm* file was constructed from the old one and filtered sequences using *cmalign*. This three-step process was repeated three times and the final *stockholm* file was the input to R-scape for visualization of the covariance model using APC-corrected G-test statistic. Alignment results across different representatives of *Pseudomonas* species were also visualized by Jalview ^36,37^.

### Phenotypic identification and quantitative PCR

Equal amount (1.0 × 10^8^ CFUs) of WT PAO1 and PAO1_ΔRsmYZ_harboring vectors with WT RsmZ and RsmZ mutants were inoculated in 200 µL of lysogen broth (LB) in sterile 96-well plate for biofilm formation, on M9-skim milk (0.5%, w/v) plate for proteolysis assay, and in 2 mL of LB for pyocyanin production. After cultured at 37°C for 24 h, the amount of biofilm was determined by crystal violet staining and quantified at OD_595_ after dissolution by 95% ethanol. Proteolysis was determined by measuring the size of proteolytic ring around the colony. Pyocyanin production was determined by extracting pyocyanin from the culture supernatant with chloroform and HCl and quantified at OD_520_. The values of each experiment were normalized to those of WT PAO1. Bacterial cells cultured in 2 mL of LB were harvested after 24 h and conducted for total RNA isolation using TRIzol (Thermo) reagents and Total RNA Isolation Kit with gDNA removal (Foregene Biotechnology). The expression levels of *lasB, rhlA, phzS, pslA* and *hcp1* were determined by quantitative PCR using an iTaq™ universal SYBR^®^ Green Supermix (Bio-Rad) and the CFX Connect Real-Time PCR Detection System with 16S rRNA as the internal reference. All the experiments were independently repeated at least three times.

## Acknowledgements

Cryo-EM data were collected at SKLB West China Cryo-EM Center and processed on Duyu High Performance Computing Center in Sichuan University. This work was supported by Ministry of Science and Technology of China (MoST) 2021YFA1301900 to Z.S., National Natural Science Foundation of China (NSFC 32070049 to Z.S., and 31970131 to K.Z.), and Sichuan University start-up funding 20822041D4057 to Z.S., J.M.B. and S.M. were supported by the Polish National Science Center (NCN 2017/26/A/NZ1/01083). Computational resources for SimRNA simulations were provided by the Poznań Supercomputing and Networking Center at the Institute of Bioorganic Chemistry, Polish Academy of Sciences through the Polish Grid Infrastructure (grant: plgsimcryox).

## Author Contributions

Z.S. conceived the project, X.J., Z.P., G.J. prepared RNAs and proteins, X.J. constructed expression plasmids, X.J, Z.P., X.L. performed binding experiments, Y.Y., X.Y., Y.W., and K.Z. carried out phenotype and qRT-PCR experiments, Y.L. collected cryo-EM data, Y.L., X.J, B.L. processed Cryo-EM data, X.J., B.L., S.M., and J.M.B. built and refined atomic models. L.L carried out covariation analysis of RsmZ secondary structure and sequence alignment. All authors contributed to the preparation of the manuscript.

## Data availability

The cryo-EM maps and associated atomic coordinate models of the complete RsmZ-A complex has been deposited in the wwPDB OneDep System under EMD accession code EMD-34048 and PDB ID code 7YR7, and the RsmZ-A complex with one missing RsmA homodimer has been deposited in the wwPDB OneDep System under EMD accession code EMD-34047 and PDB ID code 7YR6.

