## supplementary information for "Structural basis of sRNA RsmZ regulation of *Pseudomonas aeruginosa* virulence"

[#]These authors contributed equally.

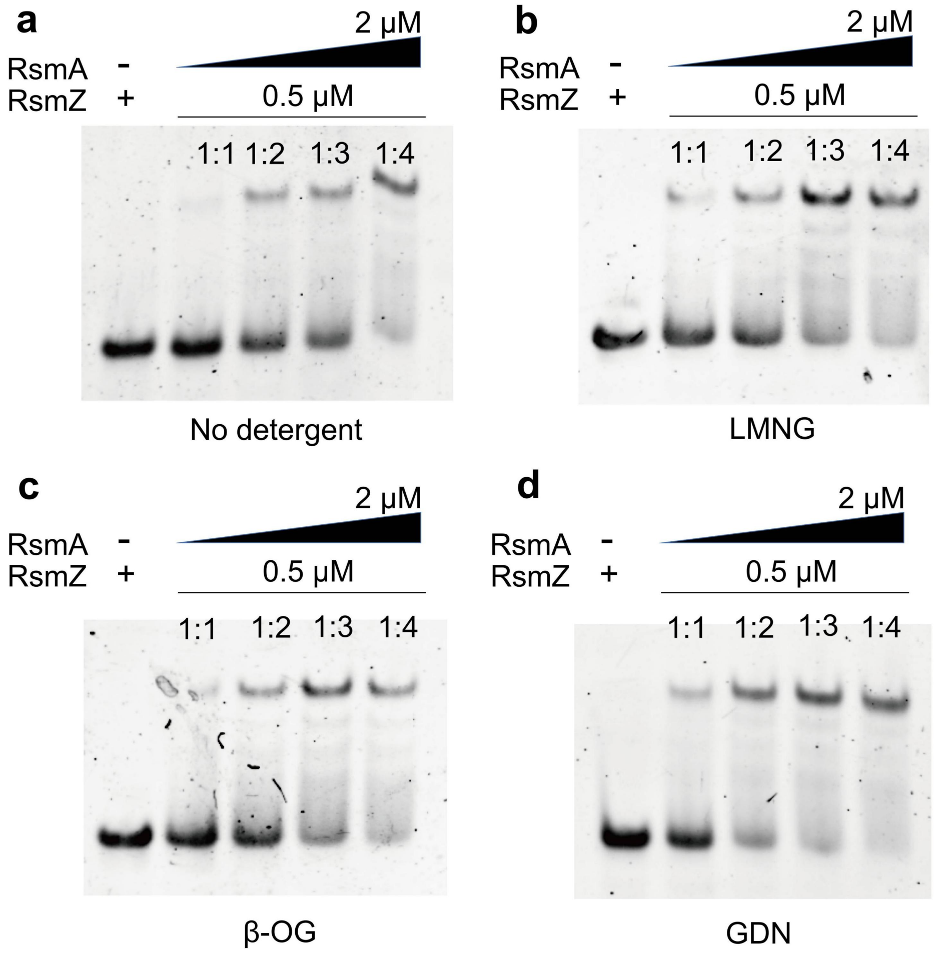


**Extended Data Fig.1 Representative EMSA analyses of RsmZ RNA complexed with RsmA protein with different detergents.** Complex formations of RsmZ and RsmA without detergent (**a**), with LMNG (lauryl maltose neopentyl glycol) (**b**) β-OG (octyl-β-d-glucoside) (**c**) and GDN (glyco-diosgenin) (**d**) were examined by EMSA.

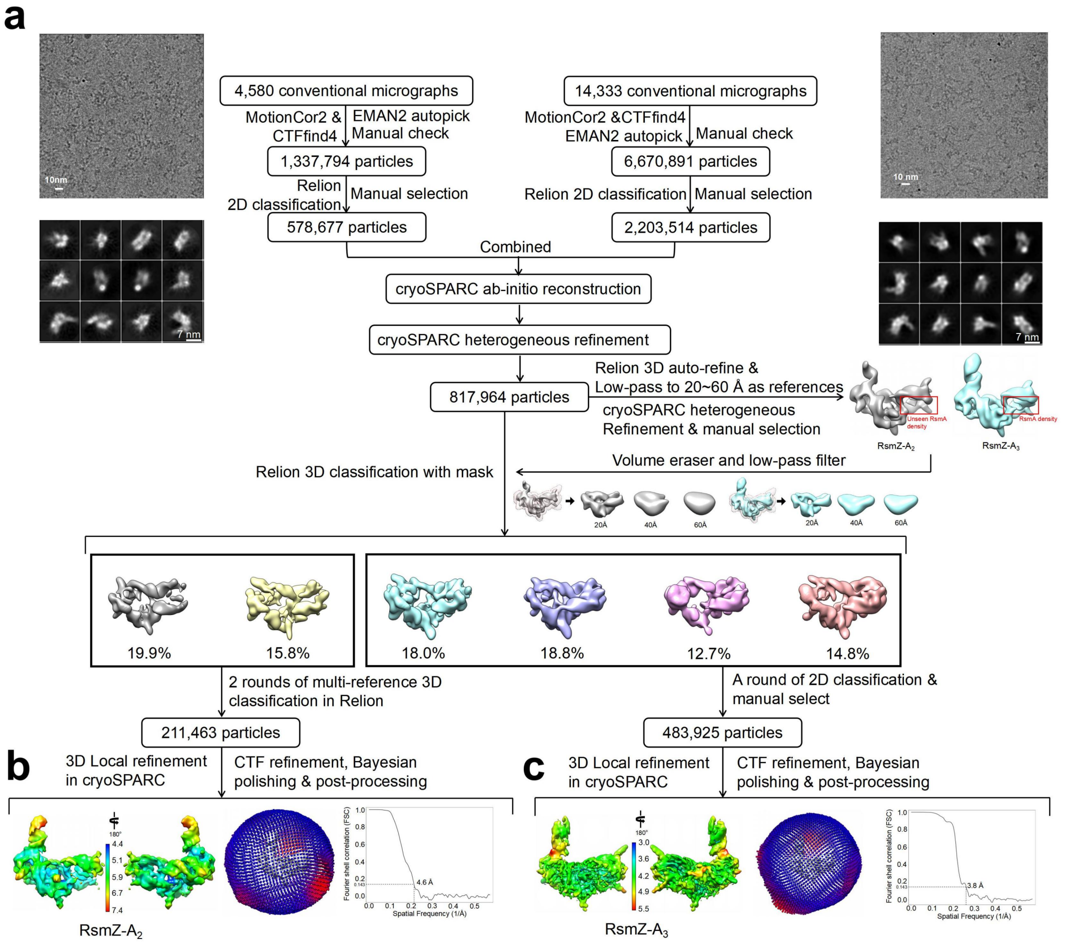


**Extended Data Fig.2 Cryo-EM single-particle reconstruction of RsmZ-A complex.** (**a**) Cryo-EM single particle workflow yielded two conformations corresponding to RsmZ-$A_2$ and RsmZ-$A_3$. Two data sets were collected from RsmZ-A complex. After 2D classification and multiple rounds of multi-reference 3D classification, the resulting particles from the smaller data set were subjected to cryoSPARC for local refinement. Red boxes indicate the RsmA density binding with SL2 and SL3 of RsmZ. (**b-c**) Colored maps according to local resolution maps with angular distribution and FSC curves indicate resolutions according to the 0.143 cutoff.

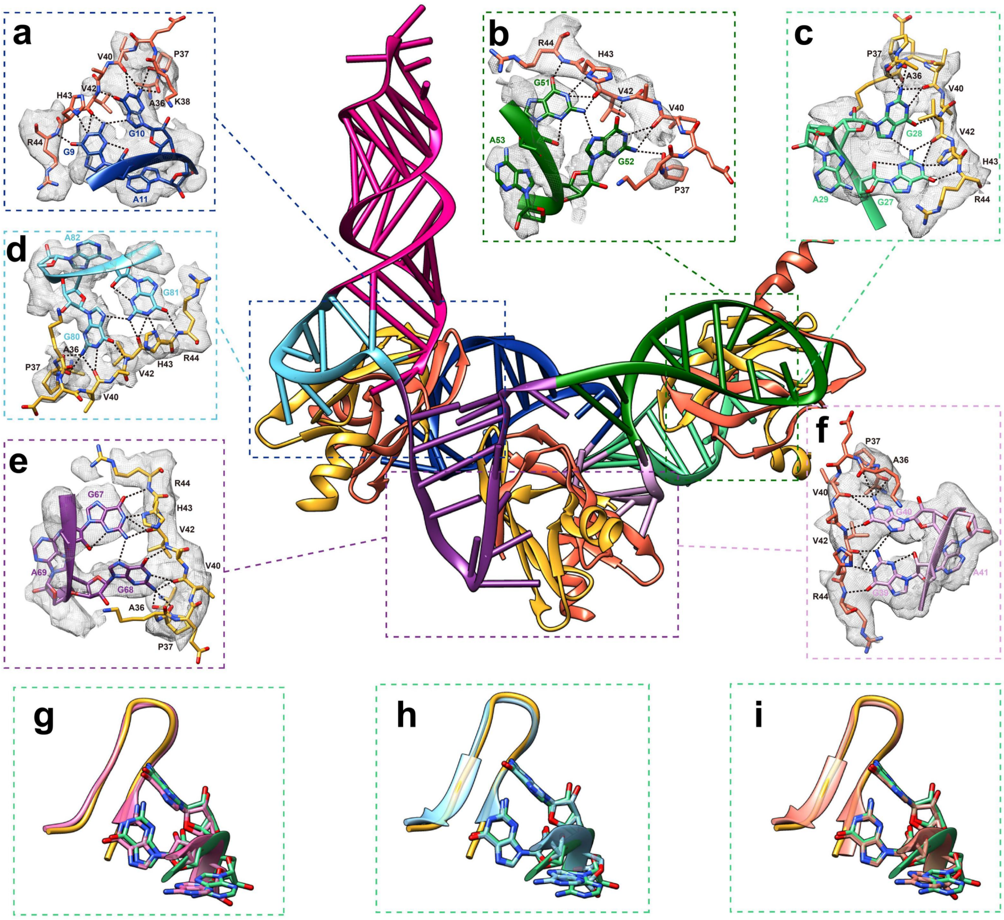


**Extended Data Fig.3 Detailed interactions between GGA binding site of RsmZ RNA and RsmA protein and superposition of *Pseudomonas aeruginosa* RsmZ-A complex cryo-EM structure with previous crystal and NMR structures of homologous complexes.** (**a-f**) Detailed interaction between *P. aeruginosa* RsmA protein (yellow and red for different polypeptide chains of one dimer) and the GGA binding site in SL1 (dark blue), SL2 (light green), SL3 (dark green), SL4 (dark purple), SL5 (light blue) and J2/3 (light purple) of RsmZ RNA. (**g-i**) Overlays of the *P. aeruginosa* RsmZ-A cryo-EM structure SL2 domain (orange) with two conformers of *P. protegens* RsmZ-E NMR structure (red, PDB 2MF0, blue, PDB 2MF1) and *P. aeruginosa* RsmN-RsmZ-2 X-ray structure (pink, PDB 4KJI).

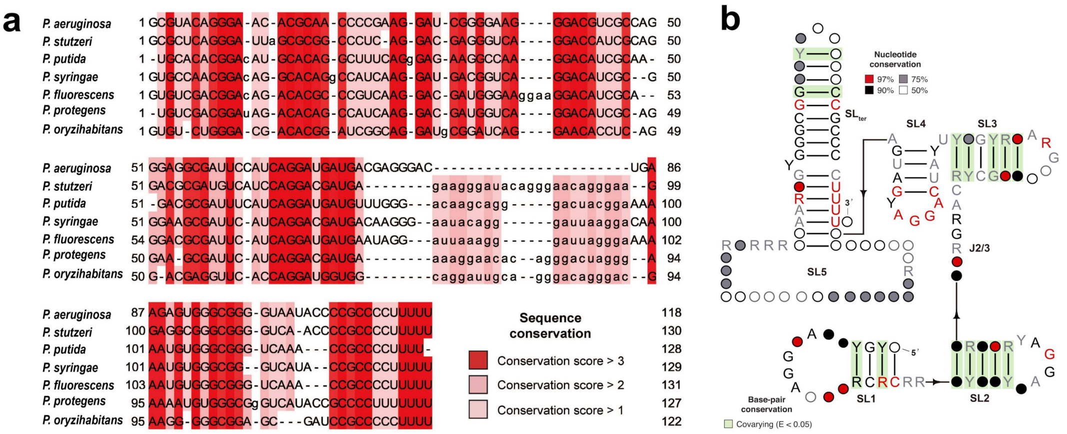


**Extended Data Fig.4 Alignment of RsmZ RNA sequences (a), and sequence covariation mapped onto the secondary structure of *P. aeruginosa* RsmZ RNA (b).** The shade levels represent degrees of conservation. Base-pairs showing significant covariation (as determined by R-scape) are boxed in green (E-value < 0.05).

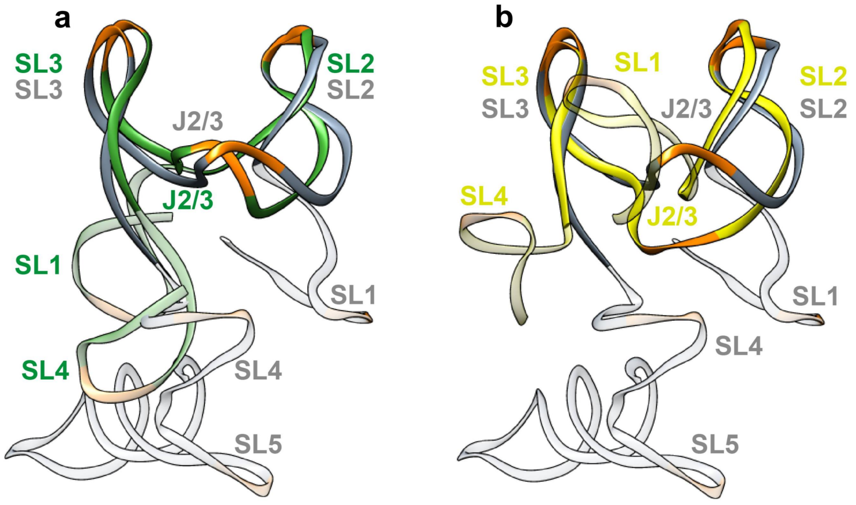


**Extended Data Fig.5 Superposition of the cryo-EM with NMR RsmZ RNA conformers.** (**a-b**) Superposition of *P. aeruginosa* RsmZ RNA structure (gray) with *P. fluorescens* RsmZ RNA conformer R NMR structure (green, PDB 2MF1) and *P. fluorescens* RsmZ RNA conformer L NMR structure (yellow, PDB 2MF0). The GGA binding sites are in orange. The different architectures were shown in a certain degree of transparency.

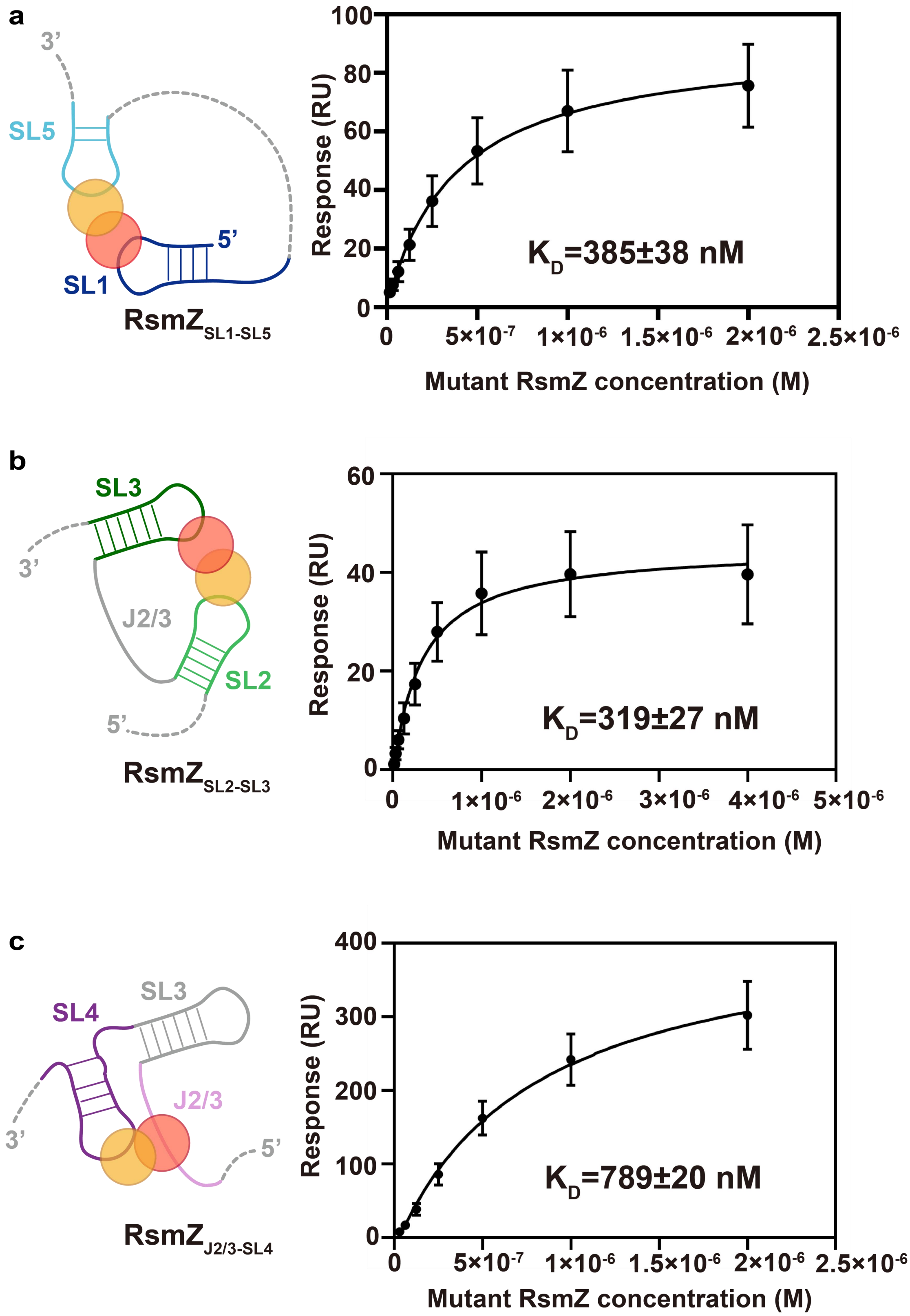


**Extended Data Fig.6 The binding curves of one RsmA homodimer in complex with different RsmZ mutants.** Binding curves and diagrams for SL1-SL5 (**a**), SL2-SL3 (**b**) and J2/3-SL4 (**c**) of RsmZ to one RsmA homodimer upon mutations of the rest of the GGA sequences are shown.
